# Data driven refinement of gene expression signatures for enrichment analysis

**DOI:** 10.1101/2024.11.03.621768

**Authors:** Alexander T. Wenzel, Farhoud Faraji, Kuniaki Sato, Kwat Medetgul-Ernar, Anthony Castanza, Romella Sagatelian, Gayathri Donepudi, Omar Halawa, Jean Y. J. Wang, J. Silvio Gutkind, Pablo Tamayo, Jill P. Mesirov

## Abstract

Gene set enrichment methods measure biological process or pathway activation in gene expression data by testing coordinate up- or down-regulation of pathway members in a ranked list of genes. These methods rely on curated, annotated gene sets whose members’ coordinate expression is an indicator of a process or state. We therefore developed the Molecular Signatures Database (MSigDB), a collection of expertly annotated gene sets. While using, enhancing, and expanding MSigDB, we have observed that some gene sets can lack coordinate expression, especially those derived from canonical pathways. To address this challenge, we developed gene set refinement (GSR), a data-driven approach leveraging large-scale multi-omics compendia to extract context-specific sets, deconvolve heterogeneity, and reveal multiple downstream signaling. We applied this method to address cancer biology questions, and demonstrated successful, targeted refinement of existing MSigDB gene sets.

## Introduction

Gene set enrichment (GSE) methods allow researchers to quantify and assess the statistical significance of changes in gene expression at the process and pathway level, providing greater interpretability than single-gene differential expression tests. Most GSE methods either test for overrepresentation of gene set genes, either in the top or bottom of a ranked list of all genes, or in a smaller set of important genes, such as the significant results of a differential expression test. In either case, GSE methods rely on the existence of pre-defined, expertly curated, annotated gene sets associated with a variety of phenotypes, processes, and pathways. A gene set intended for use with GSE should have the property that its member genes are co-expressed in samples that have the phenotype that the gene set is annotated to represent. We developed Gene Set Enrichment Analysis (GSEA), which uses a weighted version of the Kolmogorov-Smirnov statistic to test for the coordinate up- or down-regulation of a set of genes ranked by differential expression between two classes of samples [1,2]. GSEA has become a mainstay in gene expression analyses, inspiring a family of similar methods [3,4] and expansions to other types of data such as single cell assays [5].

Because GSE methods rely so strongly on high quality gene sets, we also developed the Molecular Signatures Database (MSigDB), a repository of gene sets organized into nine human and six mouse collections (www.gsea-msigdb.org) [6,7]. MSigDB includes collections such as canonical pathway databases like Reactome [8] or WikiPathways [9] (the “C2 “collection), ontologies such as Gene Ontology [10] (the “C5” collection), and cell type signatures (the “C8” collection). Each MSigDB gene set is annotated with a description of the phenotype that the gene set represents and information on provenance such as the author, the data used to derive it, and the paper in which it first appeared. MSigDB contains both human- and mouse-native gene sets. The Hallmark collection [7], released in 2015, aimed to reduce redundancy and overlap in the rapidly growing MSigDB. It was created by clustering all MSigDB gene sets by the degree of overlap in their member genes, identifying biological themes corresponding to the resulting clusters, and using them and experimental data to create and validate 50 major “Hallmark” gene sets, which have been among the most frequently accessed gene sets in MSigDB since their release.

While the Hallmarks significantly diminished redundancy among the existing collections and improved coordinate regulation in many cases, it was not as focused on the *context-specificity* of gene sets nor eradicating heterogeneity within them. By context-specificity we mean the degree to which genes are co-expressed in a specific tissue, cell type, or disease type. Gene sets may lack context-specificity for two reasons. First, they may be annotated too broadly when in fact their coordinate regulation is most evident in samples from a more specific context. This occurs most commonly for experimentally derived gene sets, which are often the top *n* differentially expressed genes in a single dataset. Second, gene sets may contain multiple context-specific components but are not coordinately regulated as a whole. This is most common in ontology gene sets that are created manually or are intended to encompass all genes related in some way to a phenotype, which is a superset of the genes that are co-expressed in cells with that phenotype.

In the last several years, the volume of data within publicly available multi-omics compendia, i.e., different modalities of data for the same set of samples, has increased substantially. We took advantage of this plethora of data and leveraged it to develop *gene set refinement* (GSR) a data-driven approach to learn patterns from across multi-omics data from several hundred samples to refine existing gene sets into one or more new, coordinately regulated gene sets. We show here that GSR can both filter a single gene set to make it more sensitive and specific by removing genes that contribute noise identifying multiple sub-signatures within that single gene set whose coordinate regulation indicates context-specific pathway activation and may yield distinct downstream signaling, enabling more robust and interpretable GSE results.

### Gene Set Refinement Overview

The GSR method combines the expression profiles for the genes in the set targeted for refinement from a chosen multi-omic compendium (e.g., DepMap [11], TCGA [12]) along with direct phenotypic measurements for the same samples (e.g., protein abundance, viability scores from CRISPR screens, etc.). It requires two inputs: a multi-omics compendium and a gene set to be refined. Appropriate multi-omics compendia contain data for several hundred to a few thousand samples. The refined gene sets are derived from the expression data (Steps 1 and 2 below) and the remaining data comprise the *direct phenotype measurements* for the gene expression datasets. Direct phenotype measurements are characteristics of samples for which enrichment of a gene set could serve as a proxy. Examples of direct phenotype measurements in multi-omics compendia include proteomics, gene dependency screens, drug vulnerability screens, epigenetic data such as methylation, mutation status, and sample metadata such as disease type. These direct phenotype measurements are used to help annotate the resulting refined gene sets (Step 3 below).

#### Step 1: Generate patterns with non-negative matrix factorization

We first define the matrix *A* as the subset of the full compendium gene expression matrix containing only the profiles of the genes in the input gene set to be refined. We then factor *A* using non-negative matrix factorization (NMF) [13]. NMF decomposes an *m* x *n* matrix into two smaller matrices *W* (*m* x *k*) and *H* (*k* x *n*) where *k* is a number of *components,* much smaller than *n* or *m,* and *A* ≈ *WH*. To avoid overfitting and to overcome NMF’s local optimality, we define consensus NMF factors using a down sampling and bootstrapping approach (see STAR Methods for details).

#### Step 2: Define new refined gene sets from NMF components

We use each component, row, of *H* to define a new refined gene set. First, we compute the correlation of each gene from *A* with that component. Unless otherwise specified, all correlations are computed with the information coefficient (IC), an information theoretic correlation metric ranging from −1 to 1 that is more sensitive to non-linear relationships [14,15]. The genes with an IC of at least 0.3 with the component comprise that component’s refined gene set. We use 0.3 as a cutoff based on benchmarking in prior work [16]. Because each component defines a new gene set, running GSR with a given parameter *k* will yield *k* new gene sets.

#### Step 3: Annotate gene sets using direct phenotype measurements

Each of the *k* resulting gene sets is characterized using its association with the direct phenotype measurements in the multi-omics data. To do this, we first score the refined gene sets using a single sample version of GSEA, ssGSEA [17], using each of the *k* refined gene sets in the test sample sets (see STAR Methods). We then compute the IC between these ssGSEA scores and the direct phenotype measurements. As we are correlating these measurements with the new gene sets’ ssGSEA scores, a strong association between a gene set and a direct phenotype measurement means that the new gene set serves as a good proxy for that phenotype when used with a gene set enrichment method.

## Results

### Refinement of ERBB2 signaling gene set yield multiple downstream signaling signatures

We applied the GSR method to the *ERBB2* signaling pathway, specifically the gene set REACTOME_SIGNALING_BY_ERBB2 [8]. *ERBB2*, the gene encoding the *HER2* protein, is a receptor tyrosine kinase which interacts with other RTKs including *ERBB3*, *ERBB4*, and *EGFR* to effect downstream changes in cell survival, proliferation, and differentiation [18]. As anti-*HER2* therapies have had success in cancers including some breast cancer subtypes, understanding context specific behaviors and mechanisms of resistance is important for improving patient outcomes. In fact, recent research has highlighted the heterogeneity of *HER2*-driven cancers and the development of resistance to anti-*HER2* treatments [19].

For gene set refinement, we used the expression data and corresponding reverse-phase protein array (RPPA) data from the Cancer Dependency Map [11], a multi-omic database of cancer cell lines. RPPA data were used as the direct phenotype measurements. GSR with k = 3, chosen by cophenetic coefficient (see **STAR Methods**), produced three distinct gene sets. Refined Set 1 yielded a gene set consisting of *GRB7, ERBB3, ERBB2, EREG, ITPR3, SRC, PRKCD, BTC, TRIB3, EGFR,* and *ADCY6*. The enrichment scores for this gene set were most closely associated with the abundance of the *HER2* protein (IC=0.50, FDR q-value=0.0001) (**Figure 2A**), indicating that this signature is more sensitive and concise signature for *ERBB2* activation than the original. Refined Set 2 yielded a gene set consisting of *NRG1, PRKCA, EGFR, MAPKAP1, SHC1, AKT3, BAD,* and *HBEGF*. In contrast, this gene set’s enrichment scores were most closely associated with the abundance of the *EGFR* protein (IC = 0.51, FDR q-value = 0.00005) (**Figure 2B**). Because *ERBB2* has context-dependent signaling activity depending on whether it homodimerizes or heterodimerizes with *EGFR* [18], these two sets emerging in different components of the refinement represent two different context-specific patterns of *HER2* signaling within the data. As an external validation for these two gene sets, we compared their enrichment scores in TCGA breast cancer samples [20] with the protein abundance of *HER2* and *EGFR* measured by the TCPA project [21]. Refined Set 1 was most associated with *HER2* protein abundance in patient tumors (IC = 0.48, FDR q-value = 0.00005) (**Figure 2D**) and Refined Set 2 was most associated with *EGFR* abundance in patient tumors (IC= 0.26, FDR q-value = 0.00005) (**Figure 2E**) as we had seen in the results using cell line data above. Finally, Refined Set 3 consists of the genes *FYN, PCG1, PRKAR2B, ADCY1, NR4A1, AKT3, PRKACA, CAMK4, RAF1, MTOR, SOS1, THEM4, AKT2, GSK3A, PRKAR1A, RNF41*, and *UBA52*. This gene set includes *PRKACA* and other components of protein kinase A. Recent research points to a role for *PRKACA* in resistance of breast tumors to anti-*HER2* treatments [22]. We therefore compared the enrichment scores for the three gene sets to the measured efficacy of various anti-*HER2* drugs measured in the same Cancer Dependency Map samples. The Refined Set 3 enrichment scores were the most anti-correlated with efficacy of Neratinib across DepMap cell lines (IC = −0.26, FDR = 0.0009) (**Figure 2C**); indicating the gene set derived from this component may represent a resistance or compensatory mechanism in *HER2*-driven tumors.

**Figure 1:**
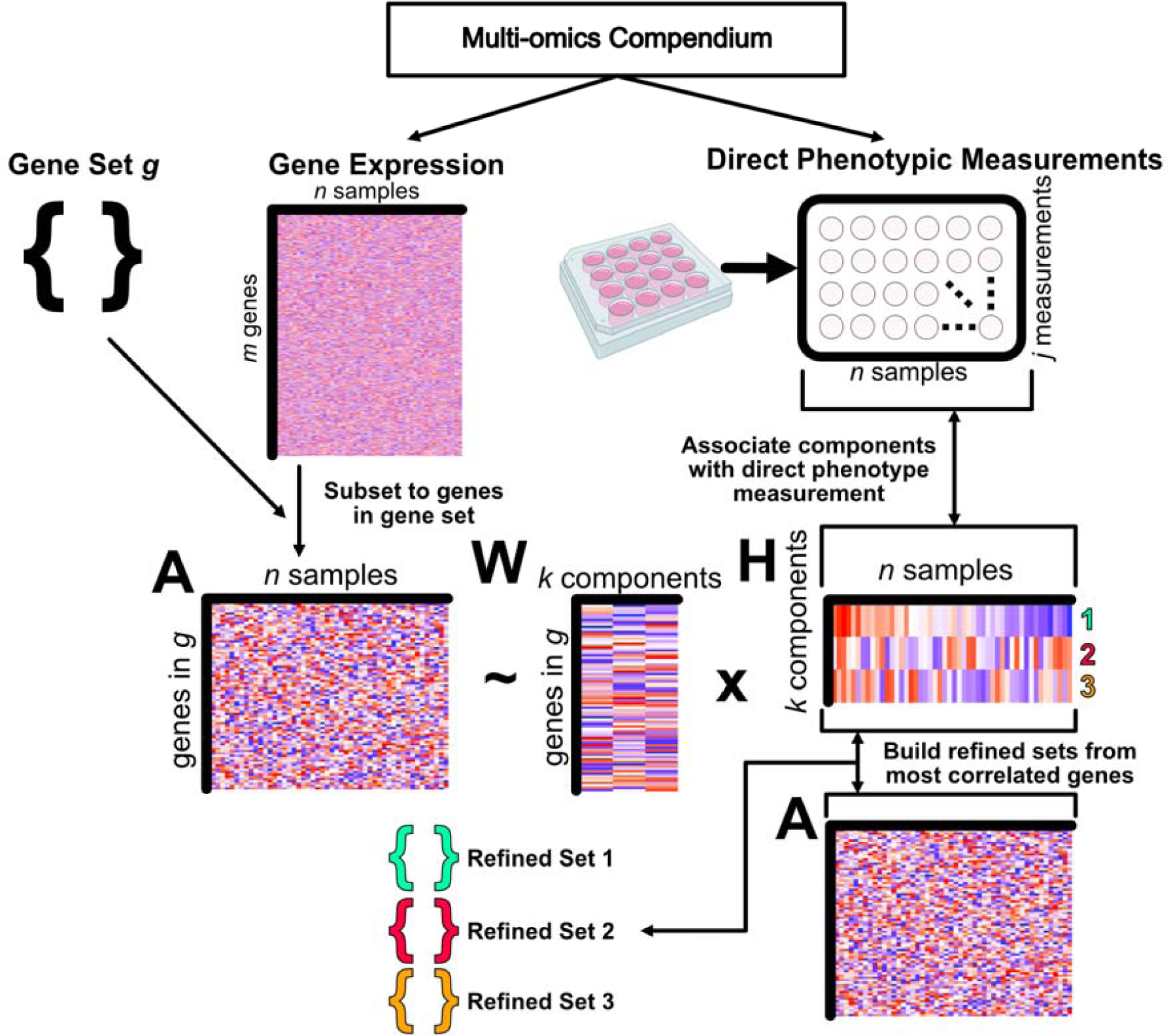
Schematic of the Gene Set Refinement method.

**Figure 2:**
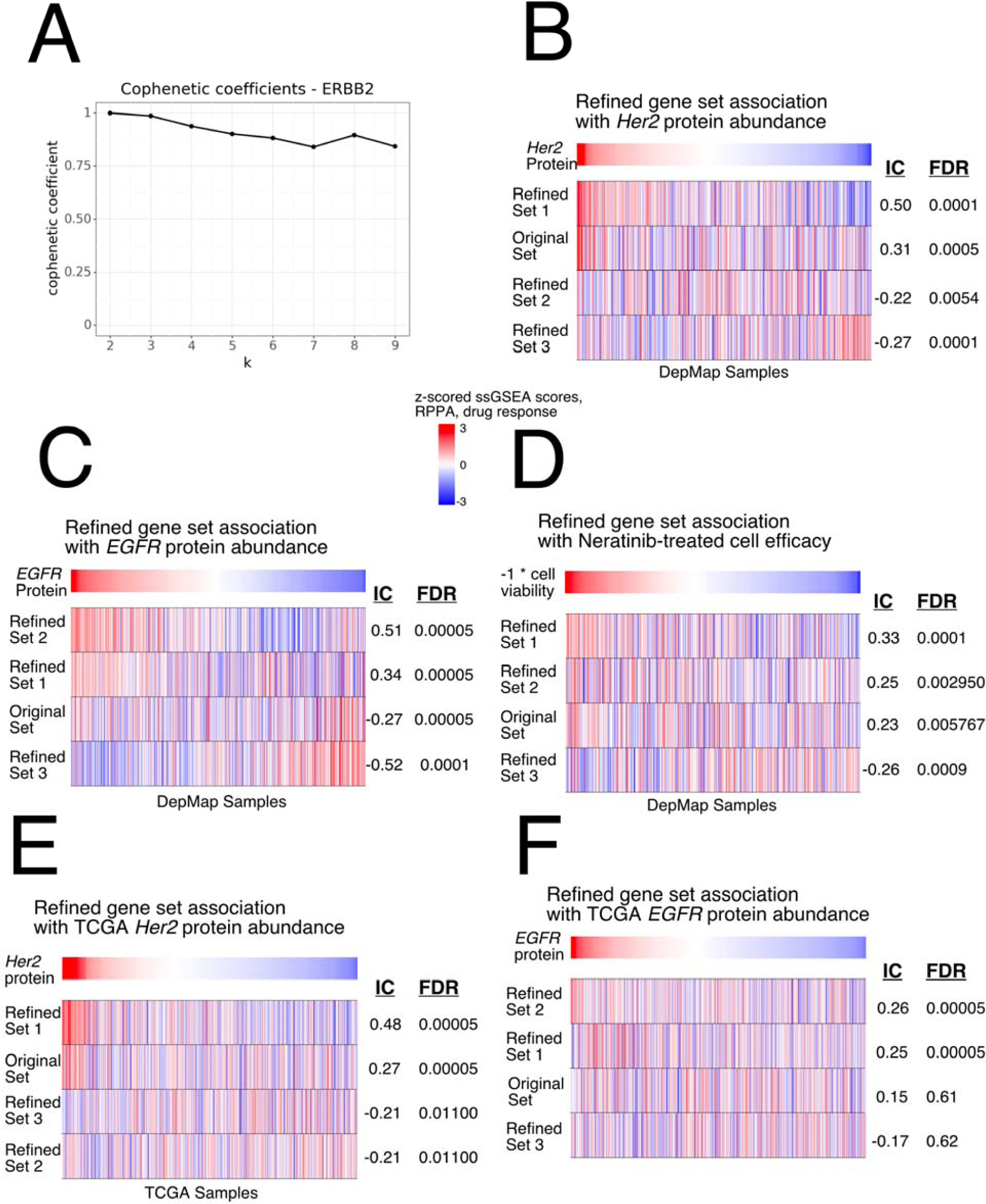
**A)** Cophenetic coefficients for the *ERBB2* signaling refinement. **B-F)** Heatmaps showing associations between ssGSEA scores and a direct phenotypic measurement in a set of samples. Top row indicates a single phenotypic measurement for the given samples, sorted from highest to lowest. Each row of the heatmap shows z-scored ssGSEA enrichment scores for a single gene set in the samples corresponding to the phenotype vector. IC: Information Coefficient. FDR is determined using a permutation test on the phenotype vector sample labels (n = 10,000). Phenotype vectors and samples used are **B)** abundance of the *HER2* protein in DepMap samples, **C)** abundance of the *EGFR* protein in DepMap samples, **D)** efficacy of neratinib as tested in DepMap samples, **E)** abundance of the *HER2* protein in selected TCGA samples, and **F)** abundance of the *EGFR* protein in selected TCGA samples.

#### Refining YAP targets reveals oncogenesis- and maintenance-specific contexts

*YAP*, along with *TAZ*, mediates transcription downstream of the Hippo pathway [23]. Nuclear *YAP/TAZ* bind with other proteins capable of binding to DNA, most frequently members of the *TEAD* family [23]. *YAP* and *TAZ* mediate transcription in response to mechanotransduction and changes in the extracellular environment [24,25] and is involved in many cancers [23,25]. Because this pathway is involved in many different cancers across cell types, we hypothesized that GSR may identify context- and cell state-dependent signatures given a list of *YAP* targets. As an input gene set, we used the validated *YAP* target genes (n=379) identified by Zanconato *et al*. [23]. GSR was run using DepMap gene expression and refined sets from k = 3 were used (**Figure 3A**), yielding three gene sets. Two of the gene sets, Refined Set 1 and Refined Set 2, were associated with abundance of the *YAP* protein (**Figure 3B**). Refined Set 1 was more associated with the abundance of receptor tyrosine kinases (RTKs) such as *EGFR*, *HER2*, and *HER3* (**Figure 3B**). RTKs are known to limit the Hippo pathway and promote *YAP*-mediated transcription via the translocation of *YAP* to the nucleus [25]. Refined Set 1 may therefore represent *YAP*’s activity as a driver of oncogenesis downstream from these genes. Refined Set 2 was more associated with proteins involved in tumor maintenance and progression. *PAI-1* (*SERPINE1*), the most associated protein, has been implicated in immunosuppression and cancer cell migration and is a marker of poor prognosis in multiple cancers [26,27]. Refined Set 2 is also associated with Caveolin-1 (*CAV1*), which has been reported to both negatively regulate EGFR as well as predict local tumor relapse and resistance to EGFR targeted therapy [28]. We therefore hypothesized that Refined Set 1 represents RTK/*YAP*-mediated oncogenesis, and that Refined Set 2 may represent *YAP*’s role in the epithelial-mesenchymal transition (EMT) and tumor maintenance. Interestingly, Refined Set 3 was anticorrelated with *YAP* protein abundance (**Figure 3B**). Because this set contains many cell cycle regulation genes such as *CCNA2* and *CDC25A*, we hypothesize that this gene set represents the behavior of genes downstream of *YAP* in non-*YAP*-dependent contexts within DepMap.

**Figure 3:**
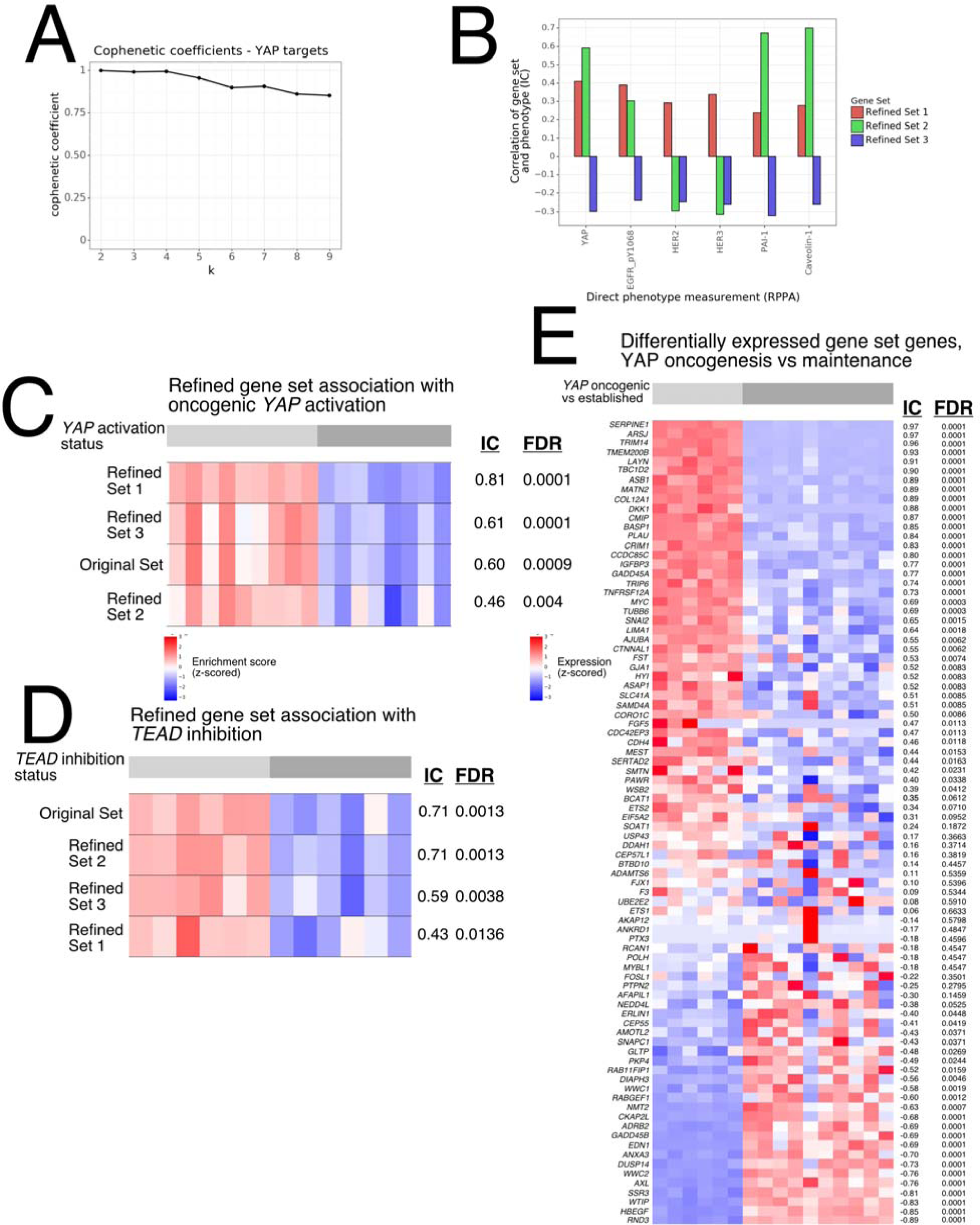
**A)** Cophenetic coefficients for the *YAP* targets refinement. **B)** Associations between protein abundances in DepMap and the corresponding ssGSEA enrichment scores for the gene sets shown. IC: Information Coefficient. **C-D)** Heatmaps with aesthetics as in **2B-F**, showing enrichment score associations with *YAP* activity in **C)** an HNSCC mouse model with constitutively active *YAP* compared to control, and **D)** HNSCC cell lines treated with a *TEAD* inhibitor or vehicle control. **E)** Heatmap of gene expression in samples from both **C** and **D**. Genes (rows) are sorted by those most expressed in the established HNSCC cell lines to those most expressed in the *YAP* oncogenesis mouse model.

We used two additional head and neck squamous cell carcinoma (HNSCC) datasets to validate the refined sets. The first models *YAP*-driven cancer initiation in which mice expressed constitutively active *YAP* and/or HPV oncoproteins. The second is a study of the effect of *TEAD* inhibition in an established HNSCC cell line Ca133. In the *YAP* oncogenesis data, we observed that Refined Set 1 best separated *YAP*-active samples from non-*YAP* active samples (IC = 0.81, FDR = 0.0001) (**Figure 3C**). In contrast, Refined Set 2 best separated wildtype samples (those with intact *YAP*-mediated transcription) from *TEAD*-inhibited samples (IC = 0.71, FDR = 0.0013). Notably, while the original gene set scored well in both datasets (IC = 0.60, 0.71 respectively), the refined sets are better able to distinguish the context specific features of these two datasets, as Refined Set 2 scored lower in the *YAP* oncogenesis dataset (IC = 0.46, FDR = 0.004) and Refined Set 1 scored lower in the *TEAD*i dataset (IC = 0.43, FDR = 0.0136).

## Discussion

We have developed a computational approach to refining gene sets for use with gene set enrichment methods. This method takes an existing gene set, and using NMF, transforms it into one or more *refined* sets, defined by NMF-derived patterns, which are more sensitive and specific indicators of a phenotype or of some contextually-specific pathway or process behavior. In the case of the Reactome *ERBB2* signaling gene set, GSR identified a decomposition into three major components. Two of these components represent two contexts of *HER2* signaling, whether homodimeric or heterodimeric with *EGFR* [18]. The third component defines a gene set that contains *PRKACA* and is anti-correlated with response to Neratinib, indicating that it may represent a compensatory or resistance mechanism to *HER2*-targeted therapy [22]. By subjecting a list of *YAP* targets to GSR, we identified two components that define two context-specific gene sets representing *YAP*-mediated oncogenesis by RTKs [25] and *YAP*-mediated tumor maintenance and progression [28,29]. Both Refined Set 1 and Refined Set 2 contained known suppressors of *YAP* such as *AJUBA*, *AMOTL2*, and *WWC1* [29], indicating that these sets are representing *YAP* activation contexts which include increased transcription of *YAP* regulators. Refined Set 1 contains *MYC* and *HBEGF* and is correlated with RTK abundance, meaning it is likely a signature of *YAP*-driven oncogenesis via RTK signaling. By contrast, Refined Set 2 contains *SNAI2*, a marker of a more EMT-like, migratory cell state [30]. Refined Set 2 also contains *AXL*, which has been shown to phosphorylate *HER3* which may facilitate cancer maintenance and progression in HNSCC [31].

The GSR method as described here has limitations which merit additional investigation. While the purpose of the method is to identify meaningful patterns in large-scale, multi-omics data for the purpose of generating refined gene sets, this does mean that the application of GSR is limited to domains of biological knowledge for which such large data exists. However, there is growing recognition of the importance of multi-omic sample characterization and data generation [32], and the utility of GSR will grow with the volume of available data. Additionally, while GSR can identify patterns without user intervention, a user must still supply a gene set that is comprehensive in that it encompasses all genes necessary for generating the patterns that define the refined set. Using the entire transcriptome as an input to refinement is not feasible both due to excessive runtime and because NMF is unlikely to identify pathway-specific nuanced signals within the space defined by a low integer value of *k*. While we will continue to use existing MSigDB gene sets as the primary source of input sets for GSR, it may also be possible to leverage protein-protein interaction networks and other systems of organizing and annotating genes and their relationships to define input sets of related genes from which GSR can derive context-specific patterns and refined sets. In conclusion, GSR computationally identifies more sensitive and context-specific gene sets for use with gene set enrichment methods and other related applications and presents the opportunity to leverage the growing volume of multi-omic data to yield new biological insights.

## Supplemental Methods

### Data Processing

#### DepMap Gene Expression

The 23Q2 release “OmicsExpressionProteinCodingGenesTPMLogp1.csv” file was downloaded and solid tumor samples were extracted. Genes with a sum of less than 5 TPM across all samples were removed. Each sample was subjected to a dense-rank normalization and scaling procedure,

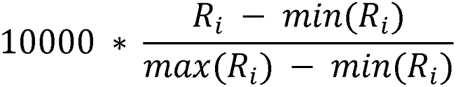

where *R_i_* is the result of applying the Python *pandas.DataFrame.rank()* function with the parameter *method = ‘dense’* using Pandas version 1.2.3.

#### DepMap Reverse Phase Protein Array

The file “CCLE_RPPA_20180123.csv” was retrieved from the DepMap Portal (release 23Q2). The data was used as originally processed [33]. Only data from solid tumor samples were used.

#### DepMap PRISM Drug Repurposing

The file “Repurposing_Public_23Q2_Extended_Primary_Data_Matrix.csv” was retrieved from the DepMap Poral (release 23Q2) and used as originally processed [33]. Each entry in the matrix is the log fold change in cell viability following treatment in cell line *i* with drug *j*. The matrix was multiplied by −1 to interpret the data as a measure of drug efficacy. Only data from solid tumor samples were used.

#### The Cancer Proteome Atlas (TCPA)

Data from the The Cancer Proteome Atlas [21] were retrieved from The Cancer Proteome Atlas Portal (https://tcpaportal.org/tcpa/download.html). Processed and batch-corrected data from “TCGA-BRCA-L4.zip” were used as provided in the portal (n = 891).

#### The Cancer Genome Atlas (TCGA)

TPM-normalized gene expression corresponding to samples available in TCPA were retrieved from the UCSC Xena portal (https://xena.ucsc.edu/). Ensembl gene IDs were converted to gene symbols using the org.Hs.eg.db R package [34]. Genes not present in the DepMap expression matrix were removed from the TCGA expression matrix.

### Gene set refinement

#### Nonnegative Matrix Factorization (NMF)

NMF was run as implemented in scikit-learn version 1.0.2 [35]. In all uses of NMF, *init* was set to “random”, *solver* was set to “mu” (multiplicative update), *beta_loss* was set to “kullback-leibler”, and *max_iter* was set to 2000. For all examples, NMF was run for values of *k* from 2 to 9 (see <u>Choosing a value of *k*</u>).

### Bootstrapping

Because sample selection and NMF random initialization can influence the final refined gene set, a bootstrapping method was implemented to prevent overfitting and generate a consensus solution. The bootstrapping approach consists of an outer and inner loop. In each iteration of the outer loop, the gene expression samples are randomly split into two-thirds and one-third. The samples in the one-third set are set aside. The inner loop is then performed on the two-thirds set. On each iteration of the inner loop, the two-thirds split is downsampled by 50%, and the remaining samples, comprising matrix *A*, are subjected to NMF, yielding matrices *W* and *H*. The pairwise correlations (using the IC) are then computed between every row of *A* and every row of *H*, which represent the gene expression and weight of the NMF component, respectively, in each sample. If *A* is an *m* x *n* matrix of *m* genes and *n* samples and NMF was run with inner dimension *k*, then the result of the entire bootstrapping process, consisting of *a* outer loop iterations and *b* inner loop iterations, is a *gene* x *component* matrix of dimension *m* genes and *a*b*k* components where an entry (*i, j*) represents the association between the *i*^th^ gene and the *j*^th^ NMF component across all NMF iterations. For all examples here, *a* = 10 and *b* = 50.

### Choosing a value of *k*

The choice of an optimal value of *k* is dependent on the expression space of a given gene set. Depending on the degree of context-specificity of complexity of the involved pathway, the different transcriptional axes may be adequately represented with a lower *k* (fewer patterns) or higher *k* (more patterns). To choose *k*, we use a heuristic that maximizes component stability and reproducibility. The *gene* x *component* matrix is subjected to NMF Consensus Clustering as described in [36]. The *gene* x *component* matrix is used instead of the *H* matrices because NMF results with different initializations are subject to random rotations [37] and different samples are used in each iteration. Briefly, NMF Consensus Clustering finds a grouping of *k* clusters using a consensus-based approach across multiple random initializations of NMF. Following a single execution of NMF, any two samples *p* and *q* are grouped together in clustering *C* if they share the same maximum component in the *H* matrix:

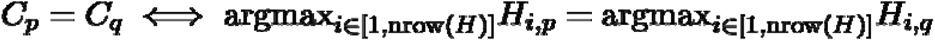

NMF Consensus Clustering builds a square connectivity matrix where entry *p, q* contains the fraction of iterations in which sample *p* is clustered with sample *q*. The connectivity matrix is then hierarchically clustered and the cophenetic coefficient, a measure of similarity between dendrogram distances and Euclidean distances in the original data [38], is calculated. This process is repeated for every value of *k*, yielding a cophenetic coefficient for each one. A high cophenetic coefficient indicates that the NMF patterns discovered at that value of *k* are distinct from one another and reproducible across sample bootstrapping and random NMF initializations. The results from the value of *k* with the highest cophenetic coefficient are used for the remainder of the method.

### Defining consensus components and creating gene sets

To define consensus patterns across bootstrapping iterations, k-means clustering is applied to the *gene* x *component* matrix. The number of clusters *k* is the same as the NMF *k* parameter. Each cluster, which consists of NMF components that have similar gene correlation profiles, defines a consensus component. Each of these consensus clusters is then used to generate a gene set. Each component consists of a number of vectors, each from a different NMF iteration, containing the correlations between each gene in the input gene set and the particular NMF component. The median of these associations is computed for each gene within each consensus component. Within each component, genes with a median association greater than 0.3, a threshold chosen based on benchmarking in previous work [16], are selected to comprise that consensus pattern’s refined gene set.

### Annotations with direct phenotype measurements

Following their creation, refined gene sets are annotated using the direct phenotype measurements. Enrichment scores are computed using ssGSEA for each of the refined gene sets in the sets of samples that were held out in the two-thirds/one-third random split in each outer loop. Within each set, the IC between the vector of ssGSEA scores and every direct phenotype measurement is computed, yielding a distribution of *a* associations between each gene set and each direct phenotype measurement. The medians of these scores are then computed and the top results for each gene set, sorted by median, are presented to the user.

## Code availability

The Python source code for gene set refinement is available in GitHub at https://github.com/alex-wenzel/PheNMF and is installed as a Python package inside a Docker image available at https://hub.docker.com/repository/docker/atwenzel/gpnb-phenmf.

